# A reagentless biosensor for mRNA: a new tool to study transcription

**DOI:** 10.1101/142794

**Authors:** Alexander Cook, Yukti Hari-Gupta, Christopher P. Toseland

## Abstract

Gene expression, catalysed by RNA polymerases, is one of the most fundamental processes in living cells. Yet, the means to study their activity are currently limited. The majority of methods to quantify mRNA are based upon initial purification of the nucleic acid. This leads to experimental inaccuracies and loss of product. Here, we describe the use of a reagentless mRNA fluorescent biosensor based upon the single stranded binding (SSB) protein. In this study, SSB showed similar binding properties to mRNA, to that of its native substrate, ssDNA. Furthermore, fluorescently labelled MDCC-SSB gave the same fluorescence response with both ssDNA and ssRNA, in a concentration dependent manner. When directly compared to RT-qPCR, we found the biosensor to be more reproducible with no product lost through purification. Therefore, the MDCC-SSB is a novel tool for comparative measurement of mRNA yield following *in vitro* transcription.

## INTRODUCTION

The information to develop and sustain life is encoded within DNA. RNA polymerases (RNAP) facilitate the distribution of this information through the transcription of DNA into mRNA. The complex activity of these machines is reflected in its large size, with a typical eukaryotic RNAPII consisting of 10-12 subunits, which provide stability, regulation and the active site for the complex (1). Transcription is a multi-stage process; the formation of the preinitiation complex, the initiation of transcription, elongation of the mRNA transcript and finally termination. All of these require multiple accessory proteins, such as the TFII family (2), CDK family (3-5) and the myosin motor proteins (6,7). Although this process has been very well characterised, yet various protein-protein interactions have still undefined roles in this procedure.

*In vitro* transcription assays have allowed the biochemical characterisation of these complex multi-protein machines. Classically, in order to measure the transcriptional activity of RNAPII, *in vitro* studies have detected the presence and/or quantified the mRNA transcripts produced. This has revealed previously unknown information about the re-initiation (8) and termination steps (9), enhancement of transcription through molecular motors (6), and has provided a method to study transcription at a single-molecule level (10).

In high yielding transcription assays, such as the T7 polymerase, the presence of mRNA can be shown by gel electrophoresis. If the analysis requires quantification of RNA, then spectroscopy is able to measure the nucleic acid concentration. However, the sample will need to be purified to remove both protein and DNA contamination, which leads to both, a loss of total RNA yield and increased experimental error. Furthermore, both approaches require a high yield of mRNA and therefore, are not typically suitable for eukaryotic transcription assays.

The use of radioactively tagged nucleotides incorporated into the transcript are common in eukaryotic *in vitro* transcription (11,12). Along with the requirement to purify the product, there are additional safety factors and costs involved. Safer alternatives do include RNA specific dyes such as the Quant-iT RiboGreen reagent (13). Reverse Transcription quantitative PCR (RT-qPCR) is a very sensitive method to quantify transcripts (14). However, contaminating DNA from the *in vitro* transcription assay can lead to large errors in the quantification. This is also a multistep process which can lead to an increase in experimental error.

Therefore, it has become clear that there is a need for a low cost, sensitive, easy to use reagent which can be added to the sample to directly compare *in vitro* transcription reactions without additional purification steps. Reagentless, fluorescence biosensors have been successfully employed in such roles in various biochemical assays (15-18).

A fluorescently labelled single stranded binding protein (SSB) from *Escherichia coli* has been successfully used as a single stranded DNA (ssDNA) biosensor for monitoring helicase activity (19-23). *E. coli* SSB is a well characterised homo-tetrameric protein containing 4 OB-fold domains (24). The formation and nucleic acid binding properties of SSB vary depending on ionic conditions. It has two main binding modes, known as (SSB)_65_ and (SSB)_35_, which have been well characterised by Lohman *et al* (25). In high salt concentrations above 200mM NaCl SSB binds to ssDNA in the form of (SSB)_65_. This case is also true when the nucleotide length is approximately 65 nucleotides long, causing the ssDNA to wrap completely around the protein. (SSB)_35_ however is a different binding mode, whereby the ssDNA wraps only around half of the protein. This binding mode occurs in low salt concentrations, less than 200mM NaCl, and when the nucleotide length is around 35 bases (26). In the case of (SSB)_35_ mode, it is possible for two ssDNA molecules 35 nucleotides long to bind to one SSB tetramer (27). SSB has been shown to have tight binding to ssDNA in both binding modes (28). Interestingly, it has been also reported to bind to its own mRNA (29).

Here, we aim to expand the application of the SSB reagentless biosensor, showing that, along with ssDNA, it can also be used to measure mRNA. This will provide a low cost, rapid and sensitive alternative for directly measuring mRNA with minimal substrate isolation. To display the functionality and versatility of the biosensor, we performed two example assays. Firstly, we directly compare the biosensor to RT-qPCR methodology for determination of transcription products. In addition, we use the biosensor to determine whether myosin VI motor activity is required during RNAPII mediated gene expression.

## MATERIALS AND METHODS

### Reagents

Unless stated otherwise, all reagents were purchased from Sigma Aldrich (Dorset, UK). Oligonucleotides are listed in Table S1.

### Protein expression and purification

SSB(G26C) was expressed from pET151 in *E.coli* BL21 DE3 cells (Invitrogen). The cells were grown to mid-log phase at 37 °C in LB media supplemented 100 μg. mL^-1^ ampicillin. IPTG was added to a final concentration of 1 mM and cells were grown overnight at 18 °C. The cells were harvested by centrifugation and re-suspended in 50 mM Tris·HCl (pH 7.5), 200 mM NaCl, 1 mM DTT, 20% Sucrose and 40 mM imidazole, supplemented with 1 mM PMSF.

For purification, the cells were lyzed by sonication and purified from the soluble fraction by affinity chromatography (HisTrap FF, GE Healthcare). The pooled protein was further purified through a Superdex 200 16/600 column (GE Healthcare) equilibrated with 50 mM Tris·HCl (pH 7.5), 1 mM DTT and 150 mM NaCl. The purest fractions were concentrated by centrifugation and stored at -80 °C.

### Labelling with MDCC

Labelling was adapted from Dillingham *et al* (20). 3 mg of SSB was incubated with 1M DTT for 20 minutes at room temperature. DTT was removed using a PD10 column (GE Healthcare, Little Chalfont, UK), equilibrated in labelling buffer (20mM Tris.HCl pH 7.5, 1mM EDTA, 500mM NaCl and 20% glycerol). After DTT removal, the solution was loaded onto the column and the column was eluted with the labelling buffer. 2-fold molar excess of MDCC (7-Diethylamino-3-((((2- maleimidyl)ethyl)amino)carbonyl)coumarin) was added and incubated for 4 hours at room temperature, with end over end mixing, while protected from light. Excess dye was removed using a PD10 column equilibrated in labelling buffer.

The concentration of SSB was taken using absorbance at 280 nm (A_280_), with extinction coefficient of *ε* = 28,500 cm^-1^ M^-1^ per monomer. MDCC concentration was determined using absorbance at 430 nm (A_430_), with extinction coefficient of *ε* = 44,800 cm^-1^ M^-1^. Labelling efficiency was calculated using equation 1.

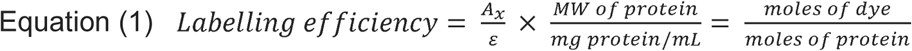

where *A*_x_ is the absorbance value of the dye at the absorption maximum wavelength and *ε* is the molar extinction coefficient of the dye at absorption maximum wavelength.

### Electrophoretic Mobility Shift Assay (EMSA)

50nM SSB was incubated with 250nM ssDNA_70_ or ssRNA_70_ for 20 minutes at room temperature in 50mM Tris.HCl pH 7.5, 100mM NaCl, 3mM MgCl_2_. Samples were loaded onto an acrylamide gel (12% acrylamide, Tris. Boric acid pH 7.5, 2.5mM Mg) (TBM) and ran in TBM buffer. SYBR®Gold (Invitrogen, Rochford, UK) stained the nucleic acids following the manufacturer’s instructions.

### Tryptophan fluorescence titration

ssDNA_70_, or ssRNA_70_, were titrated into 200nM SSB at 25°C in 50mM Tris.HCl pH 7.5, 200mM NaCl and 3mM MgCl_2_. Tryptophan fluorescence was measured using a Cary Eclipse Fluorescence Spectrophotometer (Agilent, Edinburgh, UK), with excitation at 285nm and emission at 325nm. To calculate the fluorescence quenched (%) we used equation 2.

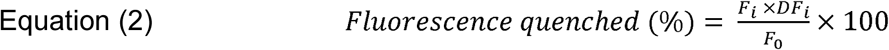

where *F_0_* is initial fluorescence intensity, *F_i_* is the intensity after titration and *DF_I_* is the dilution factor from the titration. The titration curves were fitted to equation 3:

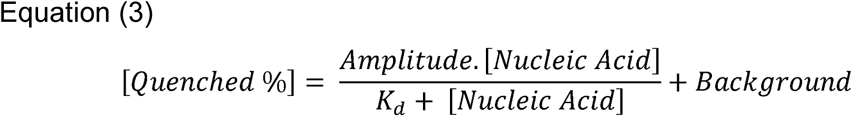

### Titrations of Oligonucleotides to MDCC-SSB

All reactions were performed at 25 °C in a buffer containing 50mM Tris.HCl pH 7.5, 3mM MgCl_2_ and 100mM, or 200mM NaCl, with 50nM MDCC-SSB in a final volume of 100 μL. Measurements were performed using a ClarioStar Plate Reader (BMG Labtech). Fluorescence excitation was measured from 400 to 440 nm, with a step-width of 1nm and emission at 470nm. Fluorescence emission was measured from 455 to 550nm, with a step width of 1nm and a fixed excitation of 430nm.The fluorescence intensity was then taken at 471nm. Fluorescence change is presented as a ratio using equation 4.

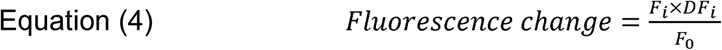

Where *F_0_* is initial fluorescence intensity at 471nm and *F_i_* is the intensity at 471nm after titration. *DF_i_* is the dilution factor for that titration. The curves were fitted to Equation 4.

### Stopped flow measurements

A HiTech SF61DX2 apparatus (TgK Scientific Ltd, Bradford-on-Avon, UK) with a mercury-xenon light source and HiTech Kinetic Studio 2 software was used. Excitation was at 435nm with emission through a 455nm cut-off filter (Schott Glass). In all experiments, the quoted concentrations are those in the mixing chamber, except when stated. All experiments were performed at 25°C in 50 mM Tris-HCl, 150 mM NaCl, 1 mM DTT, 3 mM MgCl_2_, 10% glycerol and 5μM BSA. The dead time of the stopped-flow instrument was ~2 ms: during this initial time, no change in fluorescence can be observed.

### In vitro Transcription and RT-qPCR

T7 in vitro transcription was performed using the HiScribe™ T7 High Yield RNA Synthesis Kit (New England Biolabs, Hitchin, UK) with pET28-RecD2 as a template following manufacturer’s instructions. The template was digested to yield the fragments as stated in the text. RNA polymerase II *in vitro* transcription was performed using the HeLaScribe kit (Promega). The DNA template was the pEGFPC3 linearized plasmid containing the CMV promoter which would generate a 130- base run-off transcript. Reactions were performed according to the manufacturer’s instructions. The reactions were performed for 60 min at 25°C. Reactions were also performed following pre-clearance of myosin VI from the sample using an anti-myosin VI antibody (Sigma HPA035483-100UL). Protein G Dynabeads (Invitrogen) were prepared according to manufacturer’s instructions before being loaded with 4 μg antibody. Samples were incubated for 30 min on ice and beads were extracted immediately before performing the transcription reaction. Where required, 25 μM of the myosin VI inhibitor 2,4,6-Triiodophenol (TIP) was added to the reaction mixture. RNA was then purified using RNeasy^®^ kit (Qiagen, Manchester, UK), or Gene Jet RNA purification kit (Thermo scientific), according to manufacturer’s protocol. RTqPCR was performed with one-step QuantiFast SYBR Green qPCR kit (Qiagen) using primers in Table S2. RT-qPCR samples were calibrated against known concentrations of the template.

### Cell Culture and Gene expression analysis

MCF7 cells were cultured at 37°C and 5% CO_2_, in Gibco MEM Alpha medium with GlutaMAX (no nucleosides), supplemented with 10% heat-inactivated Fetal Bovine Serum (Gibco), 100 units/ml penicillin and 100 μg/ml streptomycin (Gibco). For myosin VI inhibition experiments, MCF7 monolayers were seeded to 30-50% confluency and then subjected to 25μM TIP for 4 hours. Cells were then harvested for RT-qPCR analysis, as described above.

## RESULTS

As previously mentioned, SSB was used as a biosensor for ssDNA to report on the DNA unwinding by helicases. By building upon the work by Dillingham *et al* (20), this study used the same SSB mutant, in which the glycine residue at the 26^th^ position is replaced by a cysteine. This mutation does not affect the DNA binding of SSB, nor the formation of its tetrameric state (20). This SSBG26C mutant will be referred to as SSB throughout this study.

### SSB is an RNA binding protein

SSB has been previously reported to bind mRNA (29). However, this work was not taken any further. Moreover, the protein was reported not to bind a polyU RNA substrate (20). However, such a sequence bias may itself perturb binding and it is not representative of a transcribed gene. Therefore, before developing an SSB based mRNA, we first assessed whether SSB can bind to mRNA. To this end, we initially performed qualitative electrophoretic mobility shift assays (EMSA) with ssDNA_70_ and ssRNA_70_. Indeed, SSB bound to ssRNA_70_ in a manner indistinguishable from that of ssDNA binding (Figure 1A).

**Figure 1:**
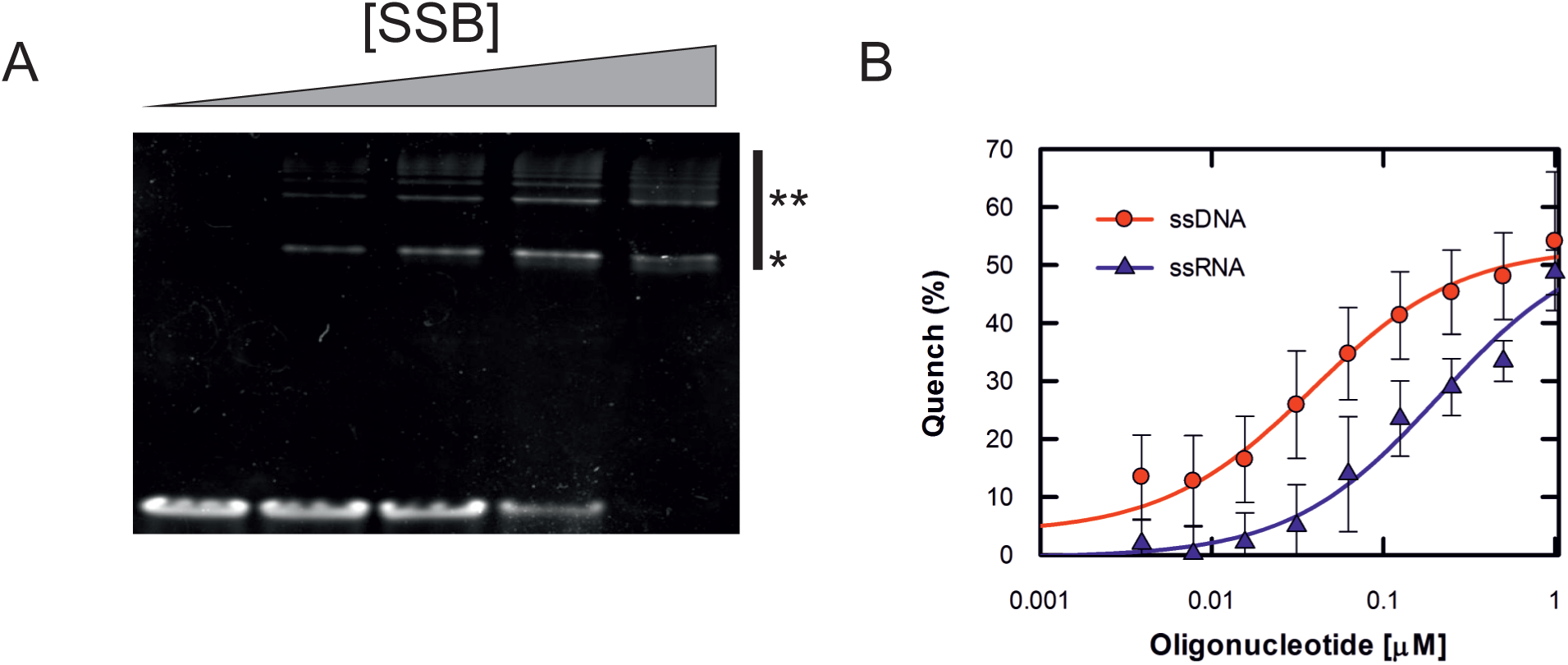
SSB can bind to single stranded RNA. (A) Representative EMSA showing qualitative association of SSB with a 70 base ribonucleotide substrate. Bound species are depicted by *. (B) Tryptophan quenching monitored while titrating ssDNA (red circles) or ssRNA (blue triangles). The curves were fitted as described in the methods. Errors bars represent SEM.

To further confirm that SSB does bind to mRNA and to define the kinetic parameters of the binding, tryptophan fluorescence quenching was used. Tryptophan residue 54 was found to be directly involved in binding to ssDNA (30), resulting in fluorescence quenching. Titration of ssDNA_70_ yielded a 50% quenching, with the apparent *K*_d_ being limited by the concentration of SSB in the reaction (Figure 1B). This was expected for the concentration MDCC-SSB and DNA used here which are well in excess of the low nanomolar *K*_d_ (31). A breakpoint in the linear phase was observed once the stoichiometric complex of 1 tetramer to ssDNA was reached. The ssRNA_70_ yielded a similar quenching response, albeit the apparent affinity was weaker (*K*_d_ ~200 nM) (Figure 1B). The weaker binding is agreement with previous findings (29), whereby SSB has preference for ssDNA over ssRNA, although binding is still possible. The consistent quenching response is indicative of a single binding site for both ssDNA and ssRNA, as would be expected from the structure.

Overall, we can conclude that ssRNA is a viable substrate for SSB and therefore, SSB is a suitable scaffold for an mRNA biosensor.

### MDCC-SSB, an RNA biosensor

To generate the biosensor, we first needed to select a fluorophore. We opted to use the commercially available and environmentally sensitive fluorophore, MDCC. This fluorophore has similar properties to IDCC which was used for the already published ssDNA biosensor (20). Previously, when IDCC was coupled to SSB through the G26C mutation, a large fluorescence enhancement of 5.3-fold was observed when SSB bound to dT_20_ (20). G26C is positioned on a flexible loop in the apo-structure, whose freedom is reduced upon DNA binding. The slight change in conformation and the DNA wrapping around the surface of SSB are thought to lead to the fluorescence enhancement. Here, the MDCC-SSB biosensor responded to the addition of both ssRNA_70_ and ssDNA_70_ substrates, as demonstrated by the fluorescence spectra (Figure 2A). A 1.9-fold increase was observed when excess ssDNA_70_ was added to MDCC-SSB, whereas a 2.1-fold increase was observed upon addition of ssRNA70. The lower fluorescence change observed here may relate to both the choice of fluorophore and buffer composition. Albeit, the signal was still sufficient for biochemical assays. Therefore, we conclude that MDCC-SSB is a suitable biosensor for ssRNA detection.

**Figure 2:**
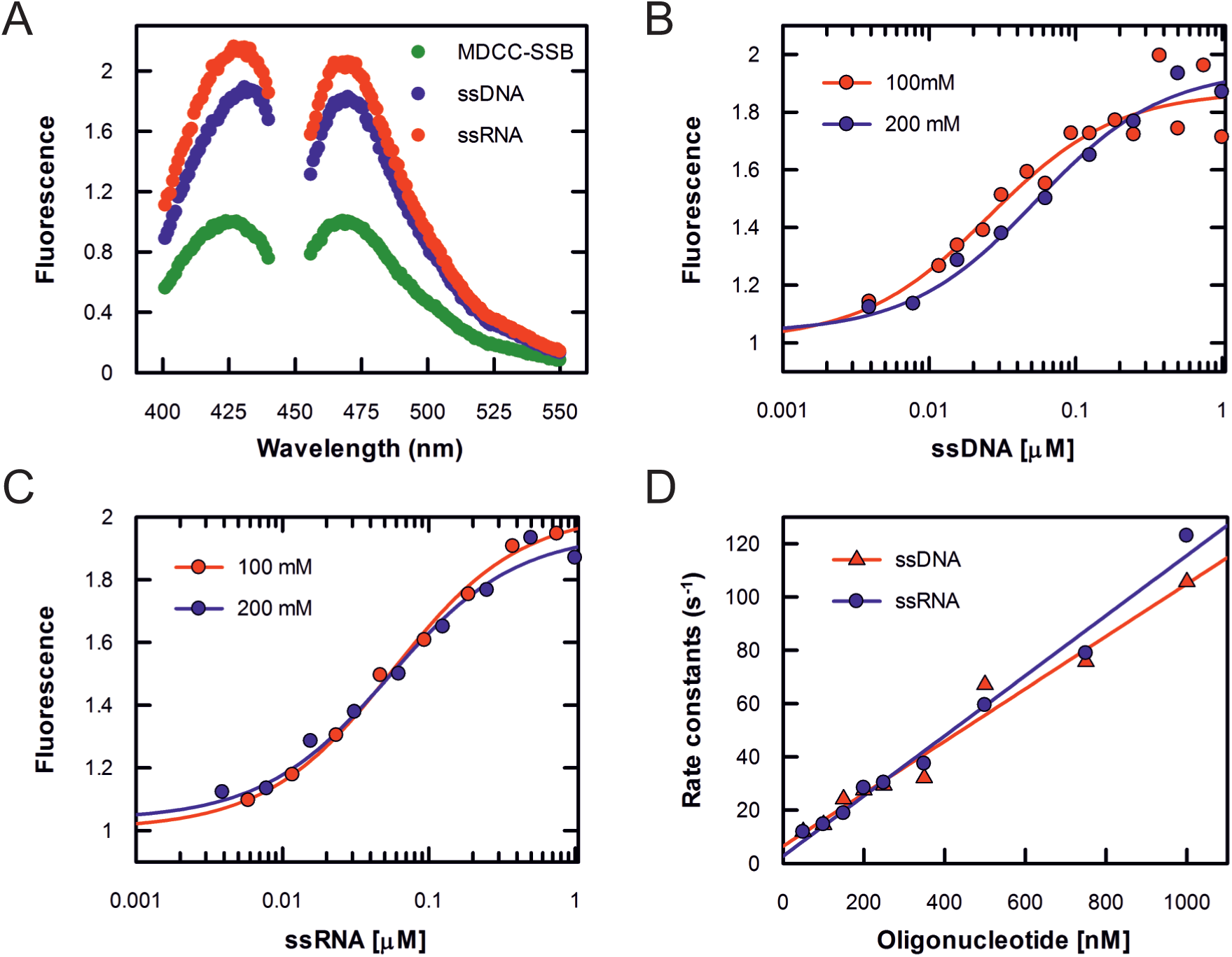
MDCC-SSB is an mRNA biosensor. (A) Fluorescence excitation and emission spectra for MDCC-SSB measured in the apo (Green), ssDNA bound (blue) and ssRNA bound (red) states. (B) MDCC-SSB fluorescence monitored while ssDNA was titrated into the biosensor in 100 mM NaCl (red) and 200 mM NaCl (blue). (C) Titration performed as in B but with ssRNA. (D) Stopped-flow pre-steady-state kinetics for MDCC-SSB binding ssDNA (red) and ssRNA (blue). Stopped-flow traces were fitted to single exponentials to yield the rate constants plotted in D. Association and dissociation rate constants were calculated from linear fits to the data. Data were averaged from three independent experiments.

### Characterisation of the MDCC-SSB biosensor

To determine the suitability of MDCC-SSB as a biosensor to quantify mRNA, we first needed to establish whether there is a dependence between the fluorescence intensity increase and the ssRNA concentration. Here, we used ssDNA as a positive control. As shown in Figure 2B, the MDCC-SSB fluorescence increase was dependent on the concentration of ssDNA_70_. As shown by the tryptophan titrations, SSB is a tight DNA binding protein (31). Therefore, any free ssDNA should be bound by the biosensor. This tight binding theoretically means that a fluorescence signal should increase linearly until a stoichiometric complex is formed, at which point the signal will be saturated. Indeed, when titrating ssDNA_70_ into 50 nM MDCCSSB tetramers, there was a clear linear phase, reaching saturation at 65 nM. This is consistent with a 1:1 complex between tetramer (and ssDNA_70_. There was a mild salt dependence on the binding affinity, whereby higher ionic strength resulted in a slightly weaker binding, which is typical of DNA binding proteins.

A similar behaviour was also observed with ssRNA_70_ (Figure 2C). The saturation point was at 101 nM MDCC-SSB tetramers, which was significantly higher than the one observed for ssDNA_70_. This binding was independent of ionic strength. The apparent weaker binding is consistent with the tryptophan titrations. Overall, the biosensor displays a large linear response to RNA concentrations, which is independent from the ionic strength at concentration between 100-200 mM NaCl.

To assess the suitability of the sensor for kinetic assays, we then measured the association kinetics of MDCC-SSB (Figure 2D). Measurements were performed under pseudo-first order conditions with a large excess of ssDNA, or ssRNA, over MDCC-SSB. For both substrates, there was a linear relationship between rate constant and concentration. The association constants were ~10^8^ M^-1^ s^-1^, which suggests rapid binding for both substrates. The association kinetics were lower than those previously published (20). The observed differences could be attributed to the presence of glycerol in our reaction buffers, which was used to stabilise the protein.

### Application of MDCC-SSB biosensor to measure in vitro transcription

We have demonstrated that the biosensor has the ability to bind to RNA 70 nucleotide long and to respond in a concentration dependent manner. To test whether the biosensor is able to also detect different lengths of mRNA at various concentrations, we used *in vitro* transcription to generate mRNA transcripts.

Transcription was driven from a T7 promoter which resulted in a 2225 nucleotide run-off transcript. Transcription assays were performed for various amounts of time, resulting in different yields of mRNA product. The product was then purified and quantified by RT-qPCR. Then, 1μM of MDCC-SSB was added to each sample of purified mRNA and the fluorescence emission spectra were recorded. Figure 3A clearly demonstrates that the biosensor was able to bind to the generated transcripts (). Moreover, there was a linear response in fluorescence intensity versus amount of mRNA, indicating that the biosensor can distinguish differences in transcript yield.

**Figure 3:**
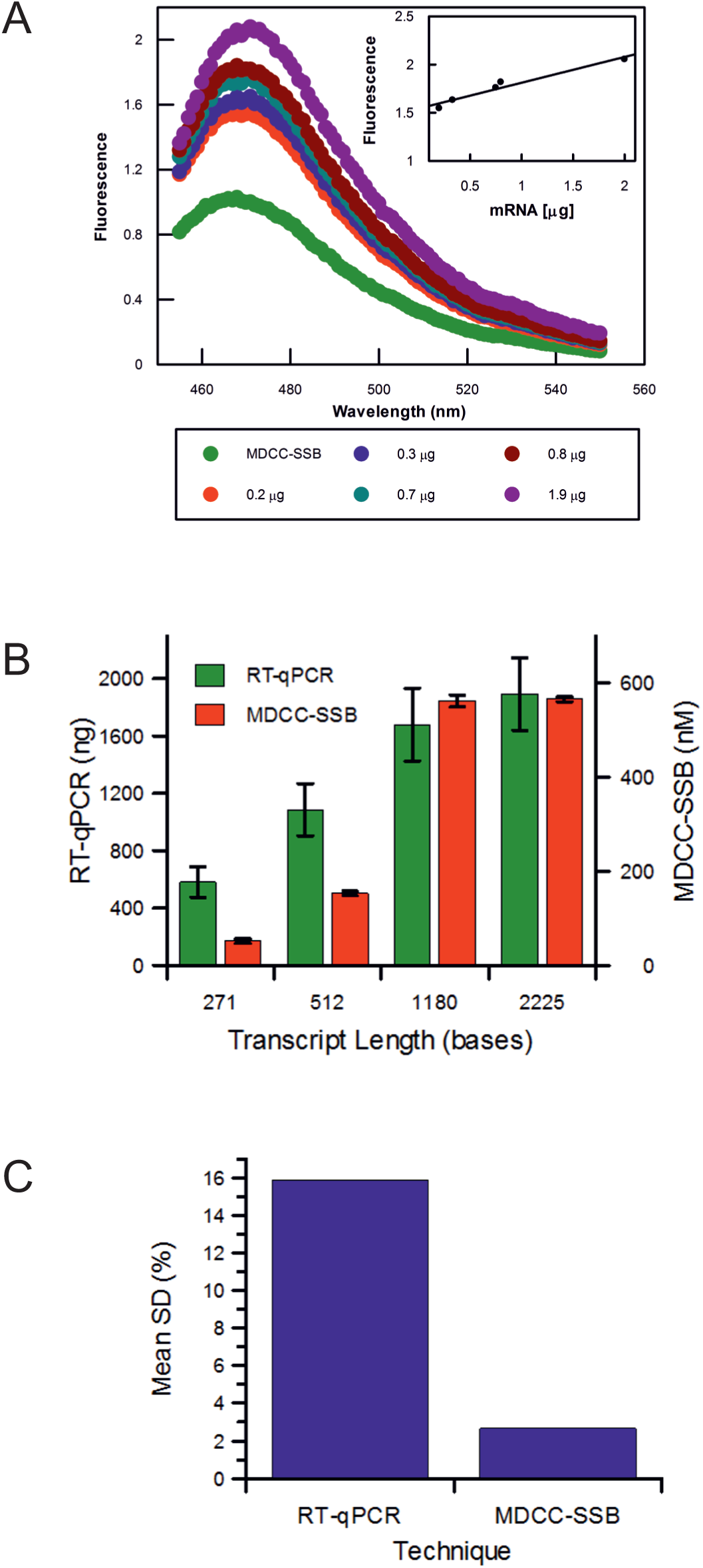
Application of MDCC-SSB to measure *in vitro* transcription. (A) Fluorescence emission spectra for MDCC-SSB and the products from 5 *in vitro* transcription assays. Experiments were performed for 20, 40, 60, 90 and 120 min then quantified using RT-qPCR using primers 1 and 2 (Table S2). (B) Comparison of RT-PCR (green) and MDCC-SSB (red) for determination of *in vitro* transcription yield. Measurements were performed as described in the Methods and text. RTqPCR and MDCC-SSB was calibrated against known substrate concentrations (Figure S1). The concentration determined by MDCC-SSB refers to nanomolar binding sites. Error bars represent standard deviation from three independent experiments. (C) Comparison between the mean standard deviation arising from the measurements in B. This is displayed as a percent of the mean value determined by the respective method.

To test whether the biosensor can be added directly into the transcription samples, without previous purification of the mRNA products, and to assess the reproducibility of the detection, we setup several *in vitro* transcription reactions to generate different lengths of transcript. At the end of the reaction, the sample was divided in two, half of which was purified and then quantified with RT-qPCR, whereas MDCC-SSB was directly added to the other half of the sample. The fluorescence intensity was recorded and was then converted into concentration of SSB binding sites (65 bases) (Figure 3B), using a calibration of MDCC-SSB against known concentration of ssRNA_70_ (Figure S1). As can be seen in Figure 3C, the RT-qPCR clearly displays larger variation in the quantification. The large differences likely result from loss of product during RNA purification and the setting-up of the PCR reaction. Most of all, it can be clearly seen that the SSB biosensor can be used to qualitatively, and quantitatively, determine differences between *in vitro* transcription assays without the need to purify mRNA.

### The MDCC-SSB biosensor as a tool to reveal the critical role of Myosin VI motor activity in gene expression

To provide an example application of the biosensor, we used it as a tool to investigate the impact of myosin VI inhibition upon RNAPII transcription. Myosin VI is critical for transcription (6), yet its role remains elusive. Performing *in vitro* transcription, combined with the rapid quantification of the transcription yield, such the one achieved with the SSB biosensor, could be a useful and efficient approach to determine the role of proteins, such as myosins, in this fundamental process.

We performed *in vitro* transcription assays using the HeLaScribe extracts. A 130-base run-off transcript was produced under the control of a CMV promoter. MDCC-SSB was added once the reactions were complete. Antibody-depletion of myosin VI has been used to perturb the activity of RNAPII (6). Our results are consistent with these findings, whereby a 60% decrease in MDCC-SSB fluorescence was observed within the depleted sample (Figure 4A). To explore whether the critical role of myosin VI involves its motor domain, we performed measurements in the presence of a large excess of F-actin, which would sequester the myosin motor domain. Indeed, this led to a 50% decrease in transcript yield. Such a result indicates a potential role of the motor activity in transcription. To further explore whether the observed perturbation was specific to myosin VI, we performed measurements in the presence of the small molecule inhibitor, TIP (32). This inhibitor has been shown to act specifically against myosin VI. Consistent with the depletion and F-actin measurements, we observed a 70% decrease in transcription. Thereby, we concluded that there is a dependence between transcription and the myosin VI motor activity.

**Figure 4:**
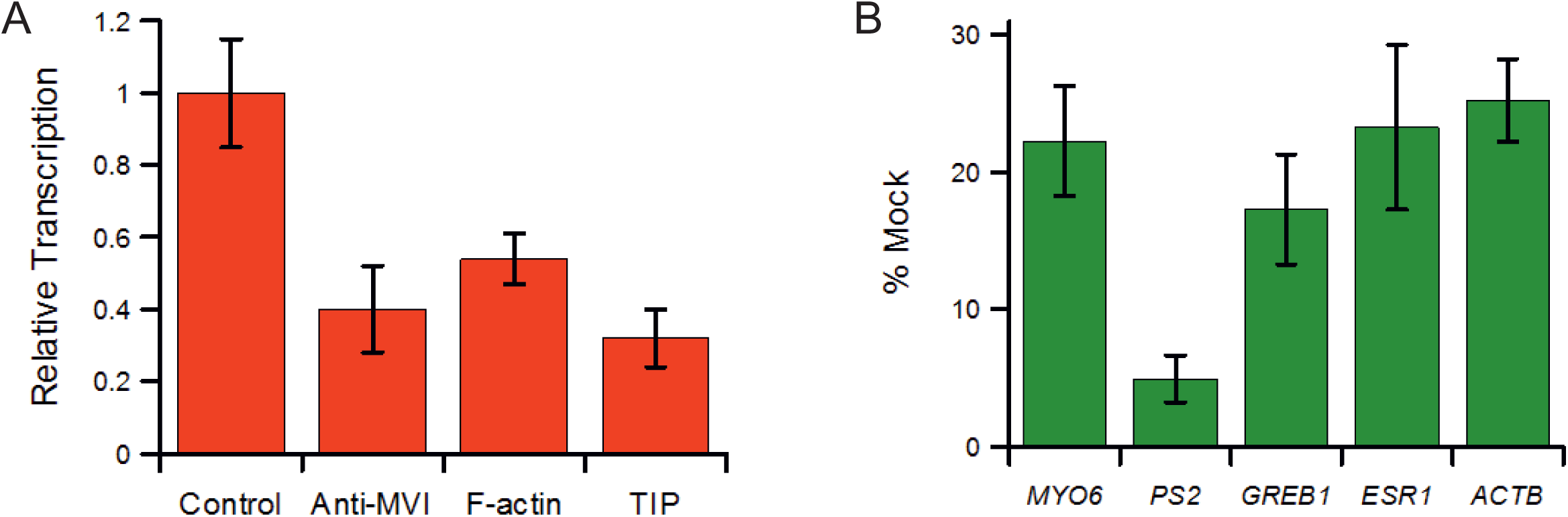
MDCC-SSB applied to study the impact of myosin VI motor activity upon RNA Polymerase II transcription. (A) *In vitro* transcription by HelaScribe extracts. Reactions were performed following antibody depletion, with the addition of 5 μM F-actin, or in the presence of 25 μM TIP myosin VI inhibitor, as described in Methods. Samples were normalized to a non-depleted control reaction (error bars represent SEM). (B) Changes in gene expression following the addition of TIP to MCF-7 cells. Expression is plotted as a percentage of expression in mock cells (Error bars represent SEM).

To understand the significance of the *in vitro* measurements upon gene expression *in vivo*, we cultured MCF-7 cells in the presence of the myosin VI inhibitor. We the monitored the expression of several genes *PS2*, *GREB1*, *ESR1* and *ACTB*. All four genes showed a significant decrease in expression (Figure 4B), indicating that myosin VI motor activity is required for gene expression. Therefore, the SSB biosensor is a quick, reliable tool which can report on changes in transcription yield, from both *in vitro* and cellular samples.

## DISCUSSION

In order to develop this biosensor, it was vital to compare the ability of SSB to bind ssRNA to its well-established ssDNA binding potential. This study has reinforced the idea that SSB is able to bind to multiple single stranded nucleic acid substrates. RNA binding has already been previously shown by Shimamoto *et al* (29), within the context of SSB’s ability to bind to its own mRNA. As DNA binding occurs through the OB-folds of SSB, it is assumed that mRNA binding would occur in a similar way, as this type of interaction does not distinguish between ssDNA and ssRNA substrates (33).

Moreover, this study showed that the addition of excess ssDNA and ssRNA results in a large fluorescence enhancement of 1.9- and 2.1-fold, respectively. The similarity in the level of fluorescence change indicates that SSB binds to both substrates in a similar way, therefore it is capable of reporting on the presence of mRNA.

The fluorescence increase occurs in a substrate concentration dependent manner for both the ssRNA and ssDNA substrates. However, the concentration at which saturation is reached is different between the two substrates. This may imply that there are differences in stoichiometry between RNA and DNA binding. For instance, the 2:1 stoichiometry of RNA:SSB complex could correspond to the 35- base binding mode. It may also indicate that there is a weaker binding affinity for ssRNA. While this would be consistent with the preferential binding to ssDNA, it remains unclear as to whether affinity or binding mode are the cause of this difference. Nevertheless, the qualitative response of MDCC-SSB to ssDNA and ssRNA is similar and the precise binding mode that underlies this response is not a requirement for utilising the biosensor. Moreover, the linear response to RNA concentration suggests that the binding is consistent and therefore, the biosensor is suitable for mRNA quantification.

The MDCC-SSB biosensor could be used in two ways: either as a qualitative comparative assay post *in vitro* transcription or as a quantitative assay following calibration. The former relies on the ability of the sensor to generate relative intensity changes between different mRNA amounts. In this case, samples can be normalised to a control and experiments can be matched accordingly. Conversely, the total mRNA concentration can be determined more accurately in terms of SSB binding sites of 65 bases. In this way, MDCC-SSB can distinguish differences between the samples based upon absolute differences in the total number of SSB binding sites, along transcripts of various lengths.

Using the MDCC-SSB biosensor offers several advantages compared to the commonly used approach of RNA purification and RT-qPCR. This is because both these two steps are time consuming and lead to a reduced transcription yield as well as large experimental errors. Conversely, the SSB biosensor can be directly be added at the end of the reaction and readily report on the transcription yield.

Moreover, this reagentless biosensor is stable for a range of salt concentrations and can be used for high-throughput analysis using a plate reader. In this way, it considerably speeds up and greatly improves the statistics of the laborious process of analysing *in vitro* transcription experiments for gene expression studies. The benefit of this approach was exemplified by investigating the dependence of transcription on the myosin VI motor activity. We found that a small molecule inhibitor of myosin VI successfully decrease transcription to a similar level to when myosin VI is depleted from the reaction. The significance of these findings in a cellular context was demonstrated by exposing mammalian cells to the inhibitor, which then led to a decreased expression of several genes tested here.

In summary, this study shows that MDCC-SSB is a reagentless biosensor suitable for measuring the concentration of mRNA following transcription. This novel tool eliminates the need for incorporation of radioactively labelled nucleotides and gel electrophoresis and increases the efficiency of the measurements through its direct use without the need of purification and RT-qPCR. In addition, it can be used to compare conditions without the need for quantification, thereby significantly speeding up the process. Furthermore, its ability to bind RNA implies that the MDCC-SSB can be used to study other biological systems including RNA helicases and potentially other RNA processing enzymes, which makes of this biosensor a powerful addition in the currently available toolbox.

## ACKNOWLEDGEMENTS

A.C thanks the University of Kent for funding his studentship. We thank Prof. M. Geeves for the F-actin sample and Dr. N. Fili for comments on the manuscript.

## AUTHOR CONTRIBUTIONS

A.C and C.P.T conceived the experiments. A.C, Y.H-G and C.P.T performed experiments and analysed data. A.C and C.P.T wrote the manuscript.

